# IMMAN: an R/Bioconductor package for Interolog protein network reconstruction, Mapping and Mining ANalysis

**DOI:** 10.1101/069104

**Authors:** Minoo Ashtinai, Payman Nickchi, Soheil Jahangiri-Tazehkand, Abdollah Safari, Mehdi Mirzaie, Mohieddin Jafari

## Abstract

IMMAN is a software for reconstructing Interolog Protein Network (IPN) by integrating several Protein-protein Interaction Networks (PPIN). Users can unify different PPINs to mine conserved common network among species. IMMAN helps to retrieve IPNs with different degrees of conservation to engage for protein function prediction analysis based on protein networks.

**Availability:** IMMAN is freely available at https://bioconductor.org/packages/IMMAN, http://profiles.bs.ipm.ir/softwares/IMMAN/.

**Supplementary information:** Supplementary data are available online.

## Introduction

Nowadays, technologies have provided access to tremendous amount of interactions at the molecular level. The study of these interactions, interactome, endeavor to model cellular and molecular processes (1, 2). Among these interactions, protein-protein interactions (PPI) are remarkable due to providing functional and structural description of executive molecules i.e. proteins. Nevertheless, PPI detection and prediction technologies are still entangling with reducing false-positive and -negative interactions (3-5). Accordingly, data integration is the best solution overall in spite of the improvement of experimental and computational methods,. STRING (6), BioNetBuilder Cytoscape app (7), IMP 2.0 (8), PINALOG (9), HIPPIE (10) and BIPS (11) are using this solution to reconstruct and refine PPI networks (PPIN). In other works, an evolutionarily conserved network with communal nodes and less false-positive links, Interolog Protein Network (IPN), was introduced as a benchmark for the evaluation of clustering algorithms (12). IPN clears up the arisen and remained interactions during evolution and help to excavate the remnants of ancestor PPIN (12-16). In this study, we present IMMAN, a software to integrate several PPINs and mine IPNs. IMMAN is free and is available as a Java program and an R/Bioconductor package.

## Methods

IMMAN enables users to define two to four arbitrarily lists of proteins (by UniProt accession number) as inputs, and seek for evolutionarily conserved interactions in the integrated PPIN or IPN as an output. Briefly speaking, the method takes the following steps to accomplish this goal.

**Step 1.** First, the amino acid sequence of each input protein list is automatically retrieved from UniProt database.

**Step 2.** In the second step, IMMAN infers the orthologous proteins. To this end, the Needleman-Wunsch algorithms is employed to compute the pairwise sequence similarities. The reciprocal best hits are retrieved and applied in the next step to increase chance of orthologous pairs discovery. The user can adjust different parameters of alignment algorithm as well as the sequence similarity cutoff for orthology detection.

**Step 3.** In this step, the nodes of the IPN are specified. Each node of the network is defined as a set of mutually orthologous proteins (OPS) such that each OPS belongs to a set of species involved in the analysis.

**Step 4.** In the fourth step, for each species, we extract singly the PPINs according to the proteins constitute the OPSs or IPN nodes. The PPINs are retrieved from STRING database. The user can adjust the minimal confidence score of STRING networks.

**Step 5.** Finally, the edges of the interolog network are computed. To this end, for every OPS pair, we count the number protein pairs (*p_jk_*, *p_jk_*) such that *p*_*i*_and *p*_*j*_ are connected in the PPIN of species *k*. If this number exceeds a predefined cutoff (coverage cutoff), there would be an edge between the aforementioned nodes. The coverage cutoff can be also adjusted by the user to tune conservedness.

## Results

After running IMMAN, the node list and the edge list of inferred IPN is produced. Additionally, IMMAN outputs the graphical representation of the network. The graphical output of IMMAN are produced using GraphViz (17) and igraph (18) in Java and R applications, respectively. The graphical representation of IMMAN on a sample dataset is depicted in Fig. 1. The sample dataset is available in Supplementary Data.

**Fig1.**
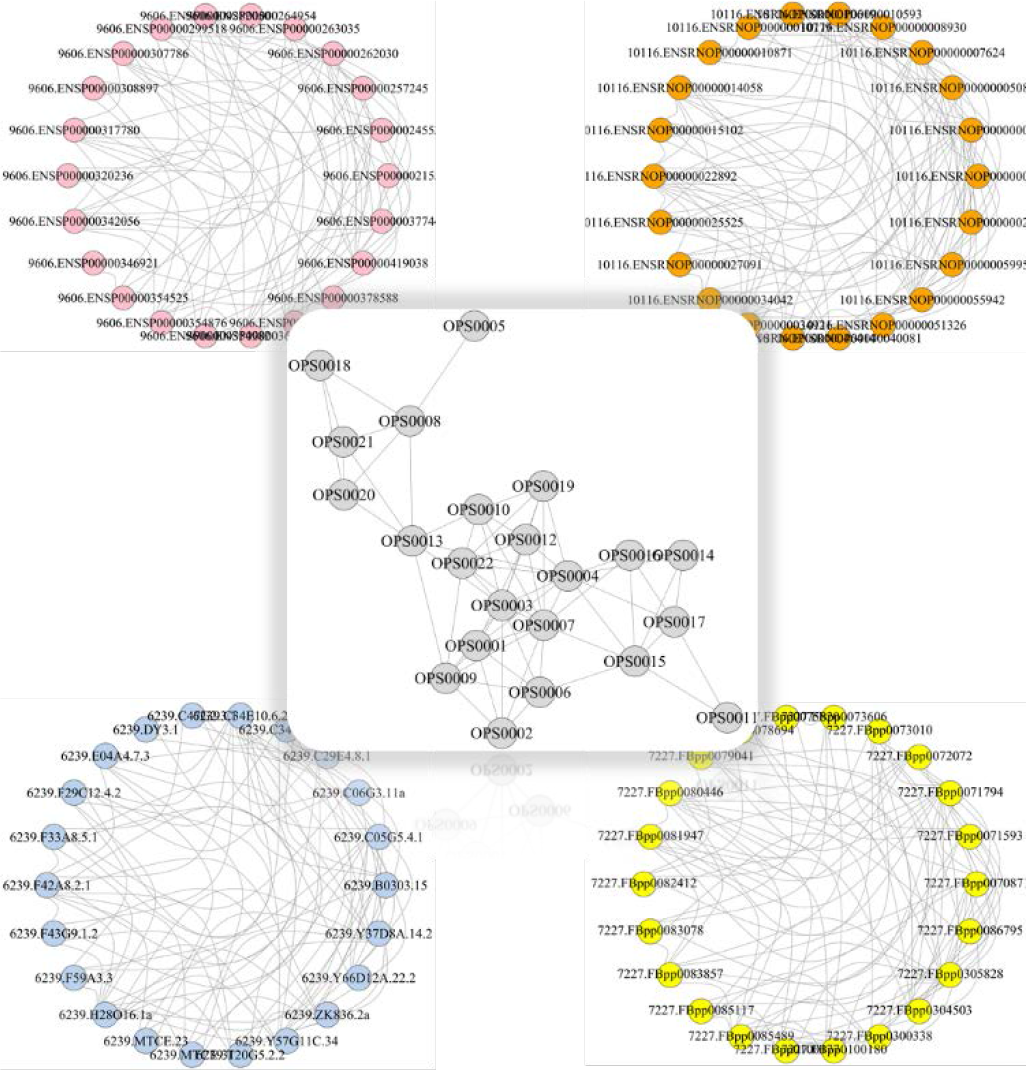
The IPN derived from four species named; *H. sapiens* (top-left), *M. musculus* (top-right), *D. melanogaster* (bottom-left) and *C.elegans* (bottom-rigth).

Although, the size of IPN is tunable by several thresholds, but obviously, missing the edges in IPN is the cost of true positive discovery which is an ideal within PPI studies with inherent inconsistency (5, 19). However, function prediction is a prominent question in molecular biology and this approach pave its way based on evolutionary mechanism (20).

## Acknowledgements

The authors would like to thank Dr. Mehdi Sadeghi for his valuable comments and discussions.

